# A Comprehensive Database of Simulations and Meshes of Coronary Arteries from the Fame 2 Trial

**DOI:** 10.64898/2026.07.30.741868

**Authors:** Fabio Marcinnò, Jochen Hinz, Edward Andò, Thabo Mahendiran, Annalisa Buffa, Simone Deparis

## Abstract

In this work, we publish the 3D unsteady Navier-Stokes numerical simulations and meshes of coronary arteries reconstructed from invasive X-ray coronary angiograms acquired during in the *Fractional Flow Reserve versus Angiography for Multivessel Evaluation 2* (FAME 2) trial. Out of the 914 clinical images, 779 vessels are successfully reconstructed and meshed. The remaining 135 vessels have been discarded since they exhibited self-intersecting geometry during the reconstruction process. The meshes are hexahedral and all of them have the same number of vertices and identical connectivity; their high quality is demonstrated using standard mesh quality indices. The simulations are performed using the Finite Element Method (FEM) with state-of-the-art coronary boundary conditions applied at the outlet. The motivation behind this effort relies from the scarcity of publicly available numerical hæmodynamics data, despite the growing interest in data-driven modeling and machine learning techniques. The database is available at the link: https://doi.org/10.7910/DVN/GPCUNS

## 1 Introduction

Coronary artery disease (CAD) is characterised by pathological changes to the coronary arteries supplying the myocardium, most commonly driven by the accumulation of atherosclerotic plaques within the arterial wall. These plaques give rise to luminal narrowings (stenoses) that progressively impair coronary blood flow. CAD manifests across a clinical spectrum. In the acute setting, it presents as an acute coronary syndrome (ACS), the most severe form of which is myocardial infarction (MI), triggered by plaque rupture and the abrupt cessation of coronary perfusion. In the chronic setting, CAD presents as chronic coronary syndrome (CCS), typically characterised by exertional chest pain (angina), where increased myocardial oxygen demand cannot be met due to flow-limiting coronary stenoses.

Invasive coronary angiography (ICA) remains the gold standard for the diagnosis of CAD. Assessment of a coronary stenosis during ICA encompasses two complementary dimensions: (i) its anatomical severity (i.e. the degree of luminal narrowing) and (ii) its haemodynamic significance (i.e. the functional impact on coronary blood flow). Beyond diagnosis, ICA also enables therapeutic intervention through percutaneous coronary intervention (PCI), most commonly stent implantation, to restore coronary perfusion.

Fractional flow reserve (FFR) is the gold standard invasive index for quantifying the haemodynamic significance of a coronary stenosis Pijls and Sels (2012); Tu et al. (2016); Achenbach et al. (2017). It is defined as the ratio of distal coronary pressure, measured beyond the stenosis, to aortic pressure, under conditions of maximal hyperaemia, typically induced pharmacologically with drugs such as adenosine Pijls and Sels (2012). Clinical studies have demonstrated that vessels with a low FFR are associated with an increased risk of adverse cardiovascular events Xaplanteris et al. (2018); Collet et al. (2026), and current European guidelines recommend revascularisation for stenoses with an FFR ≤ 0.80 Vrints et al. (2024).

However, the combination of anatomical and haemodynamic assessment is not sufficient to fully stratify MI risk. Despite being classified as haemodynamically non-significant (FFR > 0.80), up to 8% of patients with anatomically intermediate stenoses, defined as a diameter stenosis of 40–70%, still experience MI or require urgent revascularisation within two years.

Recent advances in numerical haemodynamics have made it possible to compute flow indices that cannot be directly measured during ICA. The principal numerical tools in this field, namely Computational Fluid Dynamics (CFD), Fluid–Structure Interaction (FSI), and Structural Finite Element Analysis (FEA), require the prior reconstruction of three-dimensional coronary geometries from clinical imaging modalities such as ICA and CT. Once obtained, these geometries provide a patient-specific model for simulations that enable a comprehensive analysis of coronary blood dynamics, including pure fluid flow Guerciotti et al. (2017); Menon et al. (2023) (including myocardial perfusion Papamanolis et al. (2021); Montino Pelagi et al. (2024)), blood–vessel wall interactions Malvè et al. (2012); Meza et al. (2018), and the modelling of atherosclerotic plaque progression Khan et al. (2022); Russo et al. (2023); Lissoni et al. (2025).

In the field of numerical hæmodynamics, although data-driven models have demonstrated strong potential, their effectiveness depends on the availability of large patient-specific datasets, whose scarcity remains one of the main bottlenecks limiting learning and clinical applicability Quarteroni et al. (2025). To overcome this limitation, in this work, we publish a large comprehensive database of patient-specific meshes of coronary arteries reconstructed from ICA of the FAME2 project Xaplanteris et al. (2018). For each vessel, the database contains both the result of the 3D unsteady Navier-Stokes simulation and the structured numerical mesh employed in the simulation. The numerical mesh is evaluated through standard quality indices, with the results of this assessment being reported in the database. We are able to fix the number of vertices and cells and the connectivity across all the vessels. To the best of our knowledge, in the context of numerical hæmodynamics this is the first systematic evaluation of a mesh generator conducted on such a large number of elements; we also remark that except for the SimVascular repository Updegrove et al. (2017), we are not aware of any other database that provides both simulations and meshes for real coronary arteries.

The paper is structured as follows. Section 2 provides an overview of the database and its organization. In Section 3, we describe FAME2, the dataset of invasive coronary angiographies used to reconstruct the geometrical information of each vessel. Section 4 presents the numerical meshes and discusses the results of the quality assessment. Section 5 details the model, boundary conditions, and numerical settings of the simulations. Finally, Section 6 summarizes the main motivations and conclusions behind the release of the database.

## 2 The Database Overwiew

The database is organized in the following six main directories:

- Meshes directory: This directory contains all the vessel meshes used in the numerical simulations (the meshes are scaled in meters), namely `Vessel_001.msh,…,Vessel_779.msh`.
- Quality directory: This directory holds .csv files containing the computed quality indices. For each mesh in the Meshes directory, here there is a .csv reporting the values of quality indices for each cell, namely `Vessel_001.csv,…,Vessel_779.csv`.
- `Parameters` directory:
  - `Parameter.prm`: This file outlines the parameters for the boundary conditions (for both LAD/LCx and RCA geometries), the linear solver, and all other numerical settings. Only a single file is required, as the numerical parameters remain consistent across all simulations. Simulations are performed with life^X^ Africa (2022), a finite element library for cardiac applications.
- Simulations directory. This directory contains 779 sub-directories, named `001,…779`, each representing a vessel. Each subdirectory contains:
  - The simulation results are stored in multiple files following the naming convention `solution_*.xdf5 and solution_*.h5`, where the suffix is a six-digit number that identifies each simulation instance.
  - `output.txt`: This file is the output of the solver. Details of the linear solver and time-dependent boundary conditions are reported for each time step.
  - `fluid_dynamics.txt`: This file contains the computation of some energy and turbulence related quantities, such as the total kinetic energy and the enstrophy.
  - `info.txt`: This file specifies the coronary artery type (LAD, RCA, or LCx).
- `Not-valid meshes` directory. This directory contains the 135 not valid (self-intersecting) meshes.
- `Scripts` directory. This directory includes Python scripts for boundary condition visualization and mesh quality evaluation, as well as a file identifying the 18 vessels whose simulations failed to converge. Files inside the subdirectories are not reported in Figure 1.

Both meshes and simulations can be read with Paraview Ayachit (2015). In Figure 1, the overview of the database is depicted. The database is available for download at the link: https://doi.org/10.7910/DVN/GPCUNS

## 3 The FAME2 Dataset

The dataset is composed of coronary arteries reconstructed from ICA images from the *Fractional Flow Reserve Versus Angiography in Multivessel Evaluation 2* (FAME 2) trial, a randomised controlled trial that evaluated if FFR-guided PCI would be superior to optimal medical therapy (OMT) in patients with stable CADXaplanteris et al. (2018); Collet et al. (2026). Patients with stable angina or documented silent ischemia who had at least one stenosis with a 50% diameter in a large epicardial artery that was suitable for PCI were eligible. In total, of 1220 patients, 888 had at least one pathological FFR value (*≤* 0.8) and thus underwent randomisation (447 to PCI vs. 441 to OMT). The remaining 332 patients with all FFR values >0.8 were entered into a registry, of which 166 received follow-up. For the proposed work, the 567 patients who did not undergo revascularisation were included (OMT group + registry group). Full details of this population have been reported previously Ciccarelli et al. (2018).

Of note, patient data from the FAME 2 trial were used in accordance with the original study protocol, which was approved by the institutional review board of each participating centre and conducted with written informed consent from all participants. Data were provided in de-identified form, with participants identified only by study-specific codes.

The clinical images of the FAME2 dataset comprise the three principal coronary arteries, namely the Left Anterior Descending artery (LAD), Left Circumflex artery (LCx), and Right Coronary Artery (RCA). The geometrical information, namely the centreline and radius of each vessel, is reconstructed from the clinical images using the procedure presented in Mahendiran et al. (2024). The dataset is composed of single branches (LAD, LCx, and RCA) and bifurcations are not taken into account.

Briefly, the technique relies on two ICA projections as close to orthogonal as possible. For a given ICA projection, a single frame is extracted from the recorded ICA video sequence at a fixed time point (end diastole - moment of maximal cardiac muscle relaxation). Each ICA imaging system (source and detector) is approximately placed in 3D space according to the parameters of acquisition encoded in the ICA DICOM file (primary and secondary angle, X-ray source and detector position), and the relative position of one of the two geometries is optimised in displacement to align three keypoints selected in both images. The coronary artery of interest is segmented manually from each image using a customised tablet and stylus by board-certified interventional cardiologists,Mahendiran et al. (2024), thus providing arterial thickness information (including identifying areas of stenosis) as well as the arterial centre line. The artery is described as a centreline cublic spline evolving in x and y (the pixels of the detector) as well as t (thickness in pixels). The 3D reconstruction of the centreline simply relies on the matching of spline-parameters between the two splines respecting epipolar intersections. Matching points uniquely define a 3D point in space. This is quite an easy problem for orthogonally-oriented projections since multiple intersections are rare and easy to disambiguate. The subsequent incorporation of coronary thickness information from both views allows the reconstruction of a 3D volumetric model (Figure 2). Finally, the arterial radius is reconstructed by averaging the thickness from both views, obtaining a circular cross section.

**Figure 1:**
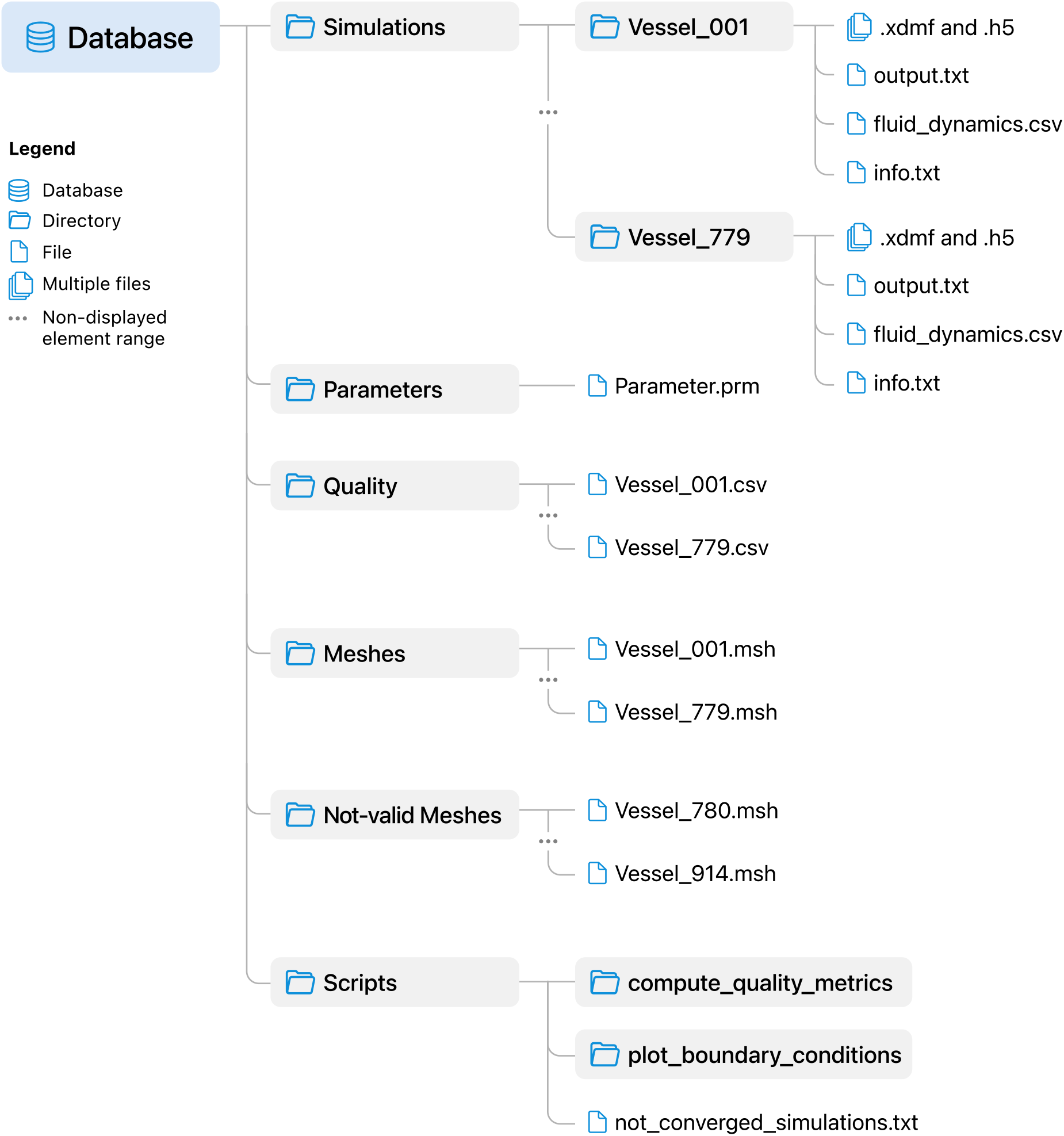
The database overview.

**Figure 2:**
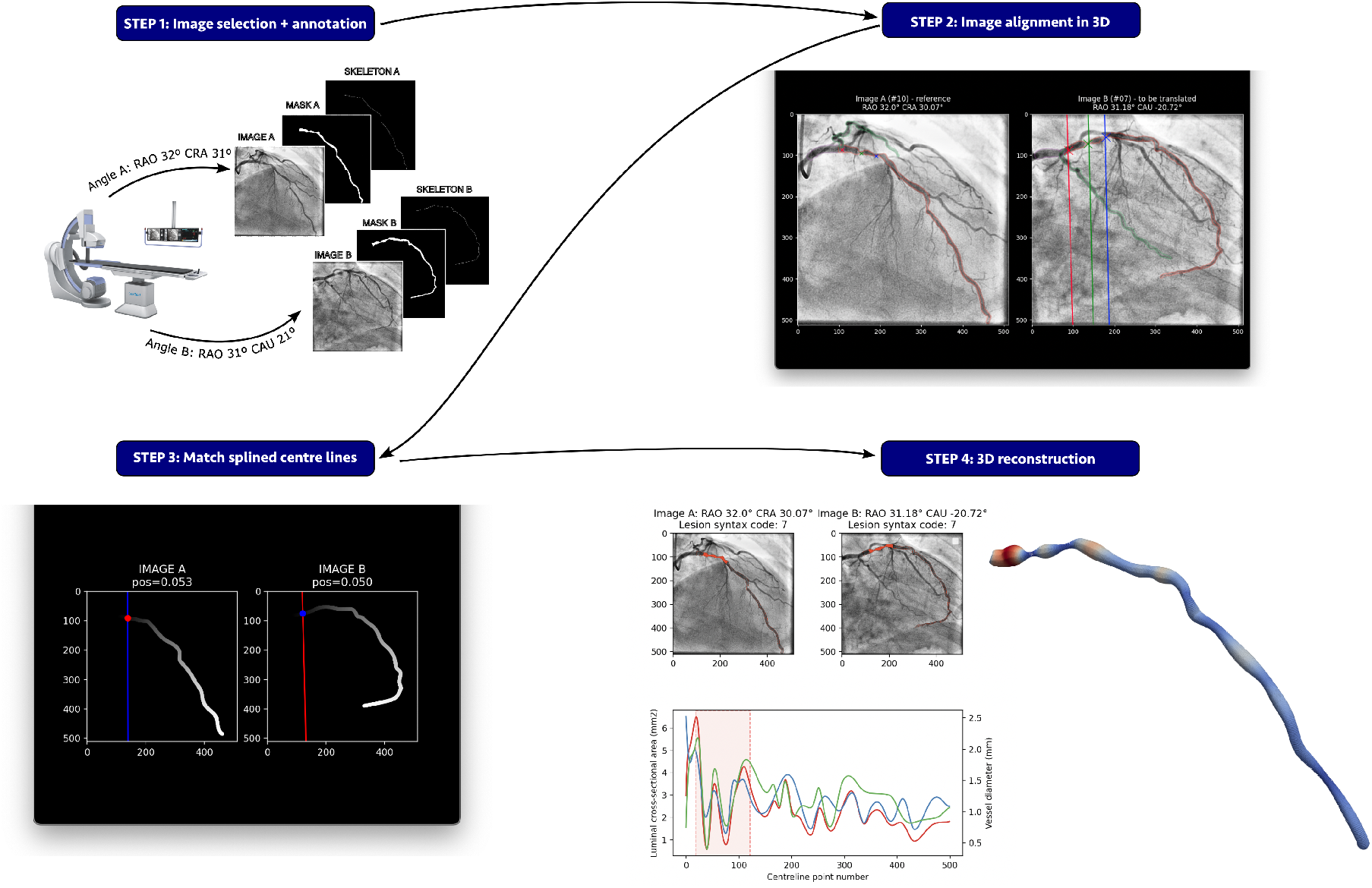
Illustrative flowchart of the reconstruction pipeline

The reconstructed geometries may contain inaccuracies arising from errors introduced during upstream processing or from violations of the underlying assumptions. Potential sources of reconstruction error include inaccurate alignment of the estimated 3D displacements, violation of the assumption that the observed motion consists solely of displacement, and inaccuracies in temporal correspondence. These errors may lead to self-intersections when the vessel surface is generated by applying the estimated thickness profile to the reconstructed 3D centreline. Such self-intersections occur primarily in regions where the vessel radius is large relative to the local radius of curvature. Of the 567 patients included in the study, ICA images were available for 561. Geometric reconstruction was unsuccessful in four cases, yielding usable data for 557 patients. In total, 914 vessel geometries were reconstructed from these patients. Of these, 135 contained self-intersections and were therefore excluded from the meshing procedure, leaving 779 vessels suitable for mesh generation. The self-intersecting geometries are nevertheless retained in the database. Self-intersections were most frequent in the RCA, occurring in 26.6% of RCA reconstructions, compared with 12.7% for the LCx and 8.3% for the LAD. This higher incidence may be related to the RCA’s characteristically tortuous, C-shaped anatomy, which makes angiographic reconstruction more susceptible to centreline overlap and foreshortening in the projected images.

In Table 1, we report the relevant statistics of the dataset.

**Table 1:**
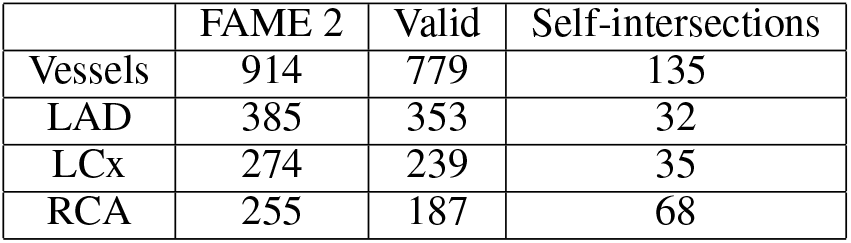
Summary of relevant dataset statistics.

## 4 Meshes and Quality Evaluation

### 4.1 Mesh generation

The structured numerical grids (`Vessel_001.msh,…,Vessel_779.msh`) are generated using the algorithm presented in Marcinnò et al. (2025).

The meshes are in the Gmsh format Geuzaine and Remacle (2009), with the inlet, outlet, surface wall, and volume regions already appropriately tagged with different labels.

It is worth mentioning that the total time needed for generating the 915 meshes, including those with self-intersections, was only of 41 minutes on a single core Intel(R) Core(TM) i7-9750H CPU @ 2.60GHz.

### 4.2 Quality assessment

Quality indices such as the Scaled Jacobian (SJ), the Edge Ratio (ER) and the Normalized Equiangular Skewness (NES) are computed for all the meshes, and reported in the files (`Vessel_001.csv,…,Vessel_779.csv`) inside the Quality directory.

All the numerical meshes have the same number of vertices and cells. Each mesh has 95’680 cells for a total of 74’534’720 cells analyzed (779 meshes analyzed). To the best of our knowledge, this is the first time that in numerical hæmodynamics a mesh generator has been systematically tested on such a large number of elements. These quality indices are computed on the hexahedra, using the ParaView filter ‘mesh quality’ Ayachit (2015), based on the vtk library Kit (2021). The code is reported inside the `Scripts` directory.

In our assessment, we use three different type of indices:

- Scaled Jacobian (SJ): Ratio between the minimum and maximum Jacobian of the element.
  - This index takes values in the interval [0, 1] and it is best practice to have such an index lie within the interval [0.5, 1] Lamata et al. (2012); Yang (2018).
  Edge Ratio (ER): Ratio between the longest and the shortest edge of the element.
  - This index takes values in the interval [1, *∞*), with a recommended range in the interval [1, 5] Yang (2018).
- Normalized Equiangular Skewness (NES): Angular deviation from ideal equiangle element configuration. It is defined as max 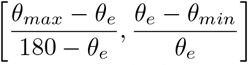where *θ*_max_, *θ*_min_ are the largest /smallest angles in the cell/face while *θ*_*e*_ is the optimal angle, which creates an equiangular face/cell (*θ*_*e*_ = 90 for a quadrilateral).
  - NES varies between [0, 1]; a value close to 1 means a degenerate cell where nodes are almost coplanar, and a value close to 0 indicates an almost orthogonal cell. The skewness should not exceed 0.5 ANSYS (2025).

In Figure 3, the histograms representing the percentage of cells within specific ranges are reported for the Scaled Jacobian, Edge Ratio and Skewness. In particular, 97.5% of the elements exhibit a SJ greater than 0.9, while 99.99% of the cells satisfy the recommended threshold of 0.5. Regarding the ER, 66.4% of the elements fall within the range 1–2, and 99.96% remain below the recommended upper limit of 5. A similar behavior is observed for the NES: 99.99% of the elements satisfy the recommended bounds, with more than 95% of the values lying within the range 0–0.2. In Table 2, we report the values of the mean and median of the quality indices, along with two additional indicative statistical measures to better visualize the quality of the elements in the extreme columns; the measure ‘% relative to median’ represents the percentage of cells in the extreme columns greater/lower (depending by the index) than the median, and the measure ‘% in rec. ranges’ states for the percentage of cells in the recommended ranges.

**Table 2:**
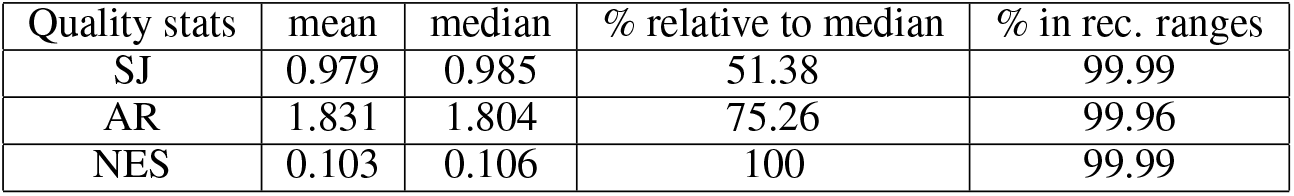
Statistics of the quality indices.

**Figure 3:**
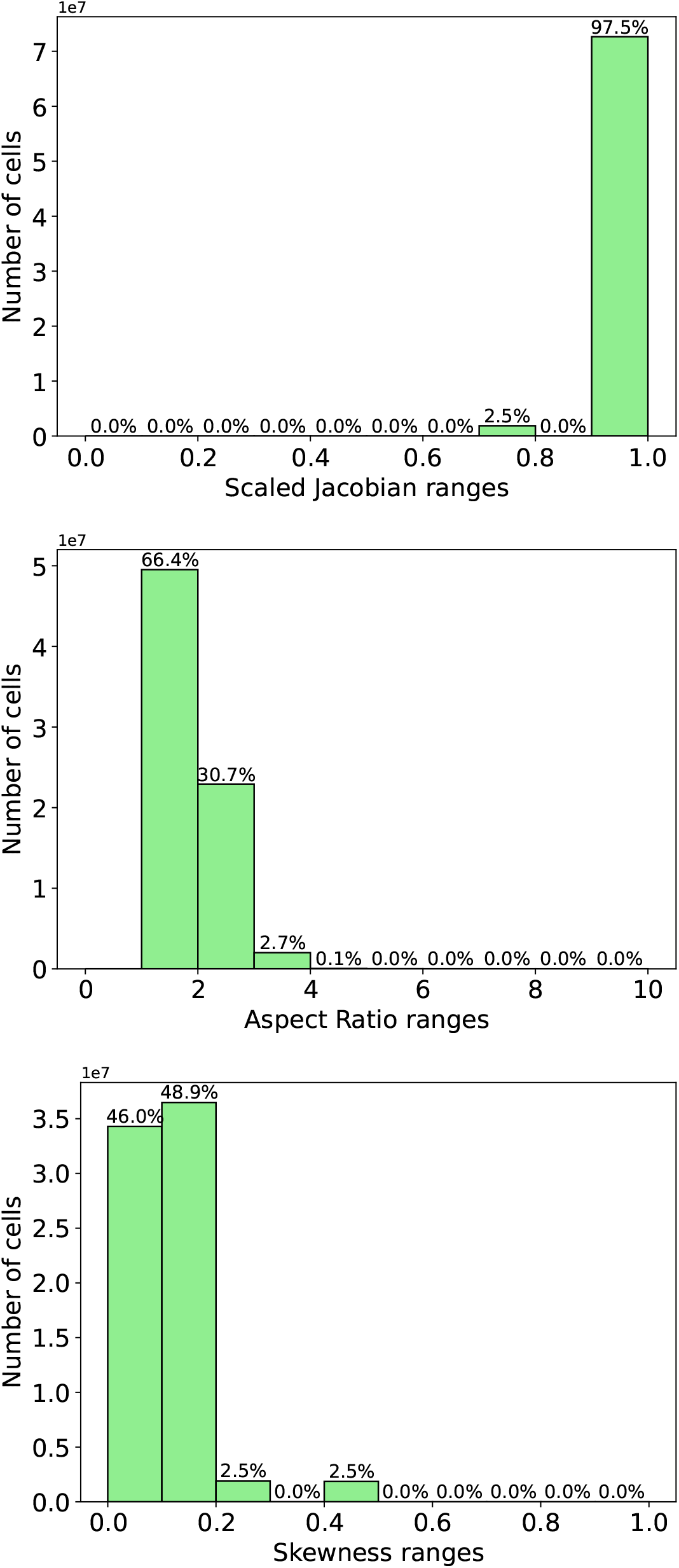
Statistics of the quality indices for the meshes used in the numerical simulations. Top: Scaled Jacobian. Middle: Edge Ratio. Bottom: Skewness.

As shown from both Figure 3 and Table 2, most of the mesh elements meet all the required standards demonstrating the high quality of the generated grids.

## 5 The Numerical Simulations

Section 5.1 describes the 3D model along with its initial/boundary conditions. This model is used for all the numerical simulations. In Section 5.2, we report the main numerical settings.

### 5.1 The model and the boundary conditions

As reported in Figure 4, we define Ω as the computational domain representing a patient-specific vessel segment, Γ_w_ as the vessel walls, while Γ_in_ is the inlet at the upstream section and Γ_out_ represents the outlet.

**Figure 4:**
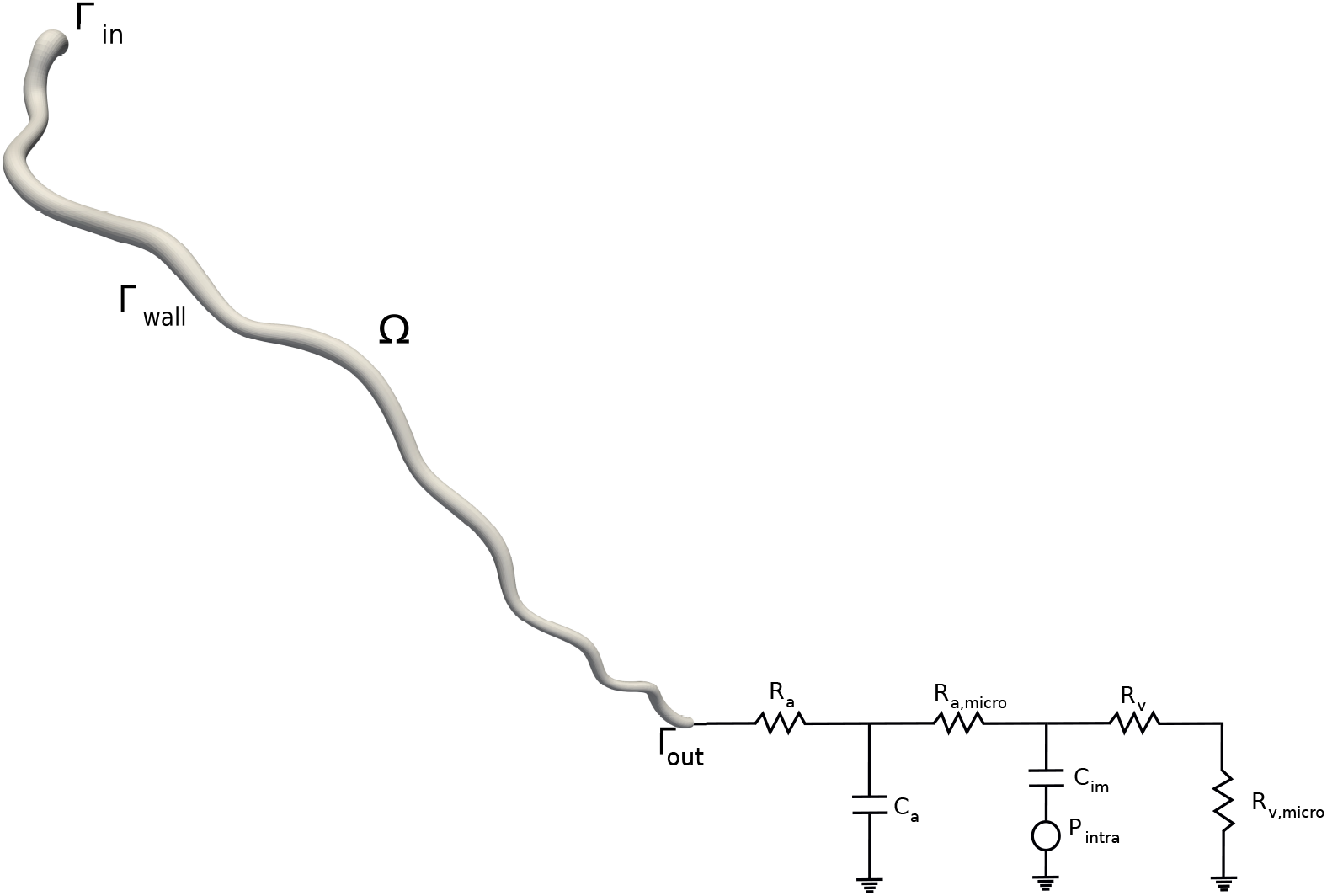
Geometric multiscale model of the coronary artery and its downstream circulation.

The fluid dynamics is modeled by means of the 3D incompressible Navier-Stokes (NS) equations that describe the blood velocity **u**(**x**, *t*) : Ω × (0, T) *→*R^3^ and the pressure *p*(**x**, *t*) : Ω × (0, T) *→* R. Defining T as the final time of the simulation, the formulation reads:Find **u**, *p* such that, for all *t* ∈ (0, 1], they satisfy:

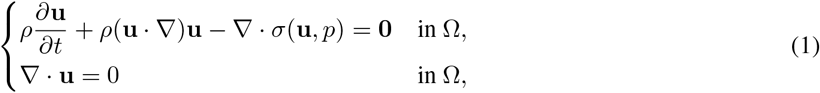

where *σ*(**u**, *p*) = *µ*(*∇***u** + *∇***u**^*T*^) − *pI* is the Cauchy fluid stress tensor, *µ* = 3.5 × 10^−3^ Pa · s denotes the constant dynamic viscosity and *ρ* = 1.06 × 10^3^ kg m^−3^ is the constant density.

Since patient-specific boundary conditions are not available, a Neumann condition (representing an aortic pressure) is imposed at the inlet section (see Figure 5, top-left). This data is generated offline by means of a closed-0D model of the entire cardiovascular system. The outlet boundary condition is imposed by means of the lumped parameter model introduced in Kim et al. (2010), see Figure 4. In Table 3, we report the values of the lumped parameters in the coronary boundary condition depending whether a LAD/LCx or RCA is simulated. In Figure 5, along with the inlet (aortic) pressure, we also report the left ventricular (pLV) and the right ventricular (pRV) pressure over two cardiac cycles used in the outlet boundary condition. All this data is reported in Parameter.prm.

**Table 3:** Lumped-parameter values used for the outlet boundary conditions, grouped by vessel type. p_LV_ denotes left-ventricular pressure and p_RV_ denotes right-ventricular pressure. The values are taken from Kim et al. (2010) and reported in SI.

| Parameter | LAD/LCx | RCA |
| --- | --- | --- |
| $R_a$ ($\text{Pa} \cdot \text{s}/\text{m}^4$) | $1.37 \times 10^{10}$ | $2.44 \times 10^{10}$ |
| $C_a$ ($\text{m}^4/\text{Pa}$) | $9.5 \times 10^{-13}$ | $2.23 \times 10^{-12}$ |
| $R_{a,\text{micro}}$ ($\text{Pa} \cdot \text{s}/\text{m}^4$) | $2.25 \times 10^{10}$ | $3.96 \times 10^{10}$ |
| $C_{\text{im}}$ ($\text{m}^4/\text{Pa}$) | $8 \times 10^{-12}$ | $1.8 \times 10^{-11}$ |
| $R_v$ ($\text{Pa} \cdot \text{s}/\text{m}^4$) | $3 \times 10^9$ | $6 \times 10^9$ |
| $R_{v,\text{micro}}$ ($\text{Pa} \cdot \text{s}/\text{m}^4$) | $4.1 \times 10^9$ | $6 \times 10^9$ |
| $P_{\text{intra}}$ (mmHg) | pLV (see Figure 5) | pRV (see Figure 5) |

**Figure 5:**
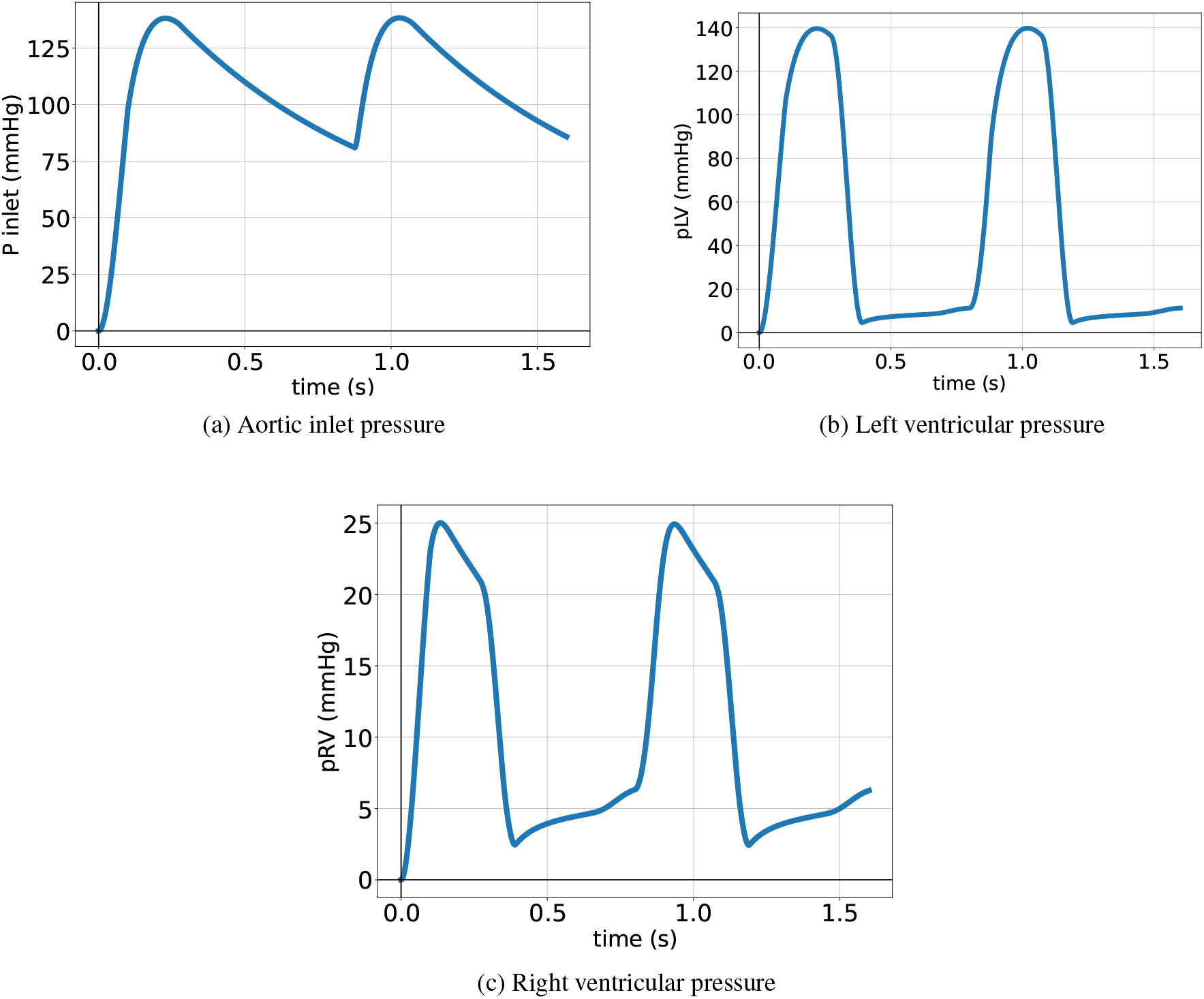
Top-Left: Aortic pressure imposed at the inlet. Top-Right: Left ventricular pressure imposed as intramy-ocardial pressure in the outlet boundary condition when a LAD/LCx is simulated. Bottom: Right ventricular pressure imposed as intramyocardial pressure in the outlet boundary condition when a RCA is simulated.

Finally, we impose a no-slip boundary condition on the vessel walls:

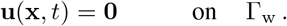

As an initial condition, we impose zero velocity and zero pressure along the entire domain.

### 5.2 The numerical settings

All the numerical parameters can be found inside the file `parameter.prm` inside the directory Parameters. For the solver details at each timestep, we refer to the file `output.txt` which is present inside every simulation directory. In this section, we briefly discuss the main parameters of the numerical simulations.

Both the 3D and 0D model equations are discretized in time using the Implicit Euler discretization, with a time step *dt* = 0.001*s*. In the Navier-Stokes equations, the non-linear term is linearized by using a semi-implicit formulation. The 3D equations are discretized in space using *Q*1− *Q*1 finite elements with the variational multiscale-streamline upwind Petrov-Galerking (VMS-SUPG) stabilization Tezduyar and Osawa (2000); Forti and Dedè (2015) and backflow stabilization Bertoglio and Caiazzo (2014) at the outlet section. The lumped parameter model at the outlet boundary is coupled with the 3D model by means of an explicit algorithm. The linear system is solved using the PETSc library Balay et al. (2022), where the system is preconditioned using SIMPLE Patankar and Spalding (1983) and the inverse of the Schur complement is approximated using the algebraic multigrid method Ruge and Stüben (1987). The linear system of 404’400 degrees of freedom is solved using the generalized minimum residual method (GMRES, Saad and Schultz (1986)) with an absolute tolerance of 10^−11^ and a forced re-orthogonalization after 5 iterations.

All simulations are performed on an Intel(R) Xeon(R) Gold 6148 CPU @ 2.40GHz with 20 cores allocated to each simulation. The library used for solving the NS equations is life^X^ Africa (2022), a high-performance object-oriented finite element library focused on the mathematical models and numerical methods for cardiac applications.

The simulations are run for 1s, where the period of the heartbeat is 0.8s. The first 0.2s should be discarded because the boundary conditions are ramped to start from zero condition (see Figure 5) to avoid spurious oscillations. We consider the first heartbeat because, with the ramping of the boundary conditions, a satisfactory regime state even within the first heartbeat is reached.

Among the 779 simulations, 18 exhibited convergence issues, i.e. the linear solver reaches the maximum number of iterations (5000). We believe these issues arise primarily due to the simplicity of these 18 geometries, mostly cylindrical with only slight curvature. In particular, such geometries appear to be more susceptible to the tuning of the outlet boundary condition parameters, leading to oscillatory behavior and convergence problems. Nevertheless, we decided to leave these simulations inside the database.

In Figure 6, we report the mesh with the velocity streamlines and pressure field zoomed on a piece of the coronary.

**Figure 6:**
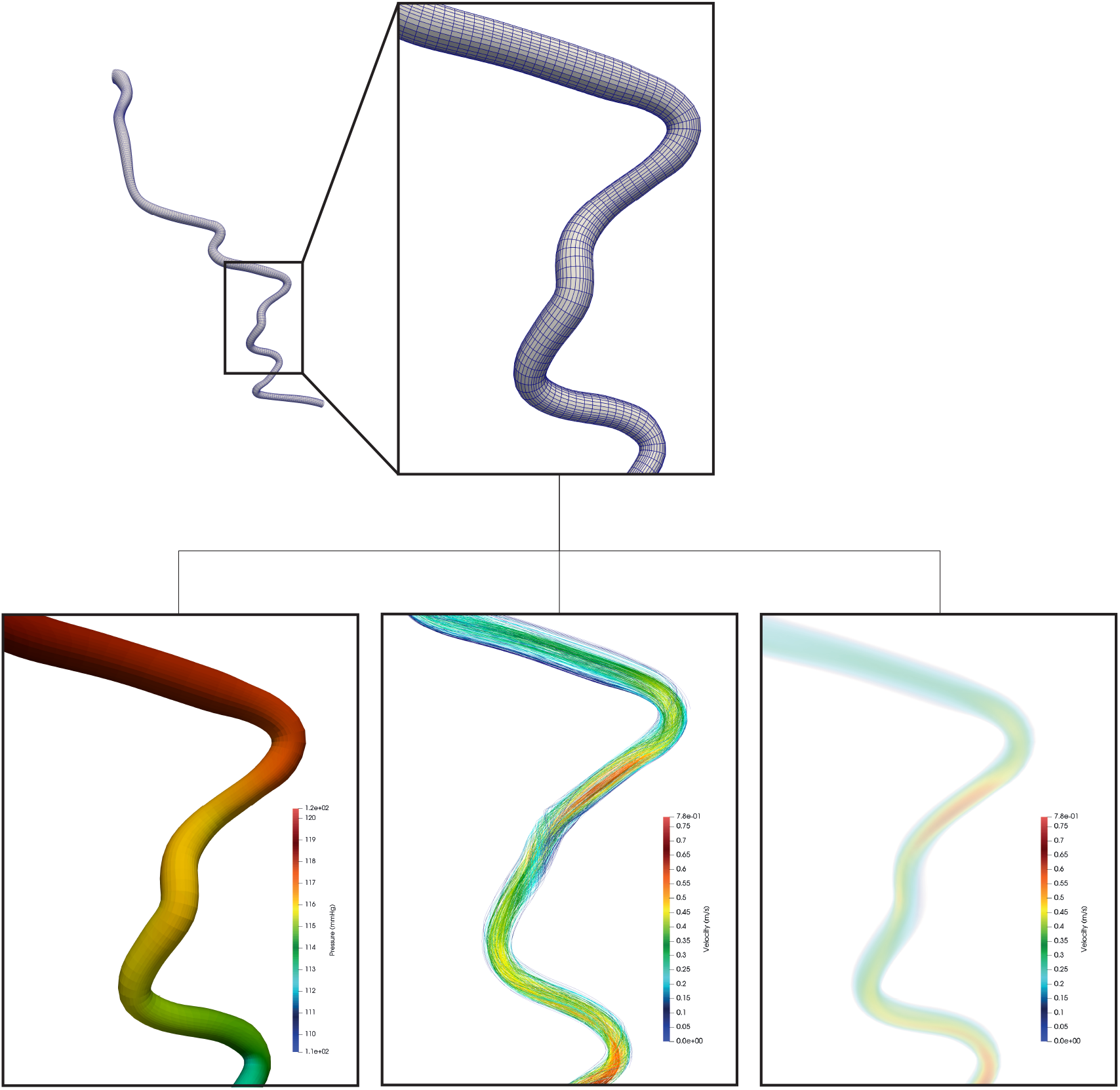
Simulation and numerical grid of a LAD artery. Pressure and velocity field along with the streamlines are depicted.

## 6 Conclusion

With this work, we release a publicly accessible database containing patient-specific coronary artery computational meshes and the results of Finite Element fluid dynamics simulations. By releasing both the meshes and the numerical simulations, we aim to give researchers the flexibility to run their own analyses, change the numerical settings, or test different boundary conditions.

We mention that a main limitation of our simulations is the use of the same values for the lumped parameters of the outlet boundary condition for each patient. While both the geometry and fluid dynamics are correctly modeled thanks to the spline-based mesh generator and the unsteady NS equations, a proper calibration of the boundary conditions should be performed, especially when a stenosis is present. This issue will be investigated in a future work.

The motivation behind this effort relies from the limited availability of open datasets in cardiovascular hæmodynamics. Despite the growing interest in data-driven modeling, machine learning techniques, and large-scale statistical analysis, the field still lacks database based on realistic anatomies. This data scarcity slows the translation of computational hæmodynamics into clinical research. We believe that releasing such a resource will support the development of more robust machine-learning models and promoting more comprehensive statistical studies in coronary hæmodynamics.

## Author contributions

F.M.: Software, Conceptualization, Validation, Methodology, Data Curation, Writing. J.H.: Writing - Review & Editing, Supervision. E.A.: Resources, Writing - Review & Editing, Data Curation. T.M.: Resources, Writing - Review & Editing, Data Curation. A.B.: Conceptualization, Resources, Writing - Review & Editing, Funding acquisition, Supervision. S.D.: Conceptualization, Resources, Writing - Review & Editing, Funding acquisition, Supervision.

## Acknowledgments

S. Deparis and F. Marcinnò have been supported by the Swiss National Science Foundation under project “Data-driven approximation of hæmodynamics by combined reduced order modeling and deep neural networks”, n. 200021-197021.

A. Buffa and J. Hinz are partially supported by the Swiss National Science Foundation project “PDE tools for analysis-aware geometry processing in simulation science”, n. 200021-215099.

Dr. T. Mahendiran (CHUV) was supported by a grant from the Swiss National Science Foundation (P500PM-210869).

Finally, we thank Dr. Bernard De Bruyne (Cardiovascular Center OLV Aalst, Belgium) for providing permission to use the FAME 2 dataset.

